# Gentle touch modulates late spinal and cortical somatosensory responses without detectable changes in spinal component N13 and nociceptive flexion reflex amplitude

**DOI:** 10.64898/2026.11.23.767102

**Authors:** A. Ribeiro-Carreira, Lídia Carvalho, Mariana Oliveira, Márcia da-Silva, Falk Eippert, Joana Coutinho, Håkan Olausson, Adriana Sampaio, Alberto J. González-Villar

## Abstract

Affective and gentle touch have been associated with changes in pain and somatosensory perception, but it remains unclear whether these effects reflect early spinal gating or later cortical and descending processes.

We investigated whether gentle touch with CT-optimal characteristics modulates spinal and cortical somatosensory and nociceptive processing across three experiments. Experiment 1 compared the sural-nerve nociceptive flexion reflex (NFR; RIII) in the leg while the forearm received no touch, vibration, robot-delivered brushing, or human-delivered stroking. Experiment 2 compared somatosensory evoked potentials (SEPs) across the same four conditions. Experiment 3 compared robot-delivered brushing with no touch, including additional trials to improve sensitivity to spinal SEP modulations.

RIII amplitudes and spinal (N13, P22) and cortical (N20, P25, N30, P45, N60, P150) SEP components were analyzed. Gentle touch modulated neither the heterosegmental NFR nor the segmental N13, arguing against inhibition of early afferent transmission within or beyond the touched segment. However, Experiment 3 revealed modulation of the P22 spinal component, suggesting that gentle touch may influence spinal processing through supraspinal feedback mechanisms. At the cortical level, both robot- and human-delivered gentle touch attenuated the N60 and P150 components relative to vibration and no-touch conditions, indicating modulation of later stages of somatosensory processing consistent with attentional and cognitive influences.

These findings suggest that CT-optimal gentle touch does not act through direct segmental inhibition of incoming sensory signals, but instead modulates later spinal and cortical processing, likely through descending influences.

This study provides a multilevel neurophysiological framework for understanding how affective touch shapes sensory processing.

## 1. Introduction

Touch is a fundamental channel through which humans connect with the world and one another, supporting bonding, social communication, and emotional development from early life [1,2]. C-tactile (CT) afferents, a class of low-threshold unmyelinated mechanoreceptive C-fibers seem crucial for mediating the affective and rewarding qualities of gentle interpersonal touch [3–5]. These fibers are found mainly in hairy skin and, although their functional anatomy remains debated, they respond preferentially to hair deflection and to stroking or brushing at the velocities and temperatures characteristic of human caresses, with responses often linked to subjective ratings of pleasantness [6–8].

Affective and gentle touch, whether CT-targeted or not, has been linked to pain regulation and modulation of somatosensory processing at behavioral and cortical levels [9–12]. However, it remains unclear whether gentle touch, possibly mediated by CT afferents, modulates somatosensory or nociceptive processing as early as the spinal level, and where along the pathway these effects first emerge.

Animal studies indicate that wide dynamic range (WDR) neurons in Rexed lamina I respond to both CT and nociceptive inputs, providing a potential substrate for interactions between gentle touch and nociceptive transmission [13]. CT-related signals then ascend toward the posterior insula, involved in interoceptive, affective, and nociceptive processing [14,15].

In humans, spinal and cortical somatosensory processing can be examined using somatosensory evoked potentials (SEPs), while the nociceptive flexion reflex (NFR) provides an index of spinal nociceptive excitability. SEPs elicited by peripheral nerve stimulation follow the fast Aβ-mediated dorsal column–lemniscal pathway and can be recorded at successive levels of the neuraxis, allowing modulation to be localized to spinal or cortical processing stages [16–18].

The NFR (RIII reflex), in contrast, is a polysynaptic spinal withdrawal reflex elicited by noxious electrical stimulation, mediated by Aδ-fiber input to the dorsal horn and producing flexor muscle contraction [19,20]. Because it is strongly shaped by inhibitory interneuronal circuits, it provides a direct and objective marker of spinal excitability and nociceptive inhibition [19,21–23]. Also at the spinal level, the N13 component reflects segmental postsynaptic activity in the cervical dorsal horn [24,25], and has been proposed as a marker of dorsal horn function and central sensitization [26,27]. Its latency may also be sensitive to social context, with shorter latencies reported for touch delivered by another person than for self-touch [28]. The later P22 component is less well characterized. Although described as a spinal postsynaptic response associated with afferent depolarization [29], recent evidence suggests that it may also receive supraspinal influences and be sensitive to attention [16]. Together, N13 and P22 provide complementary indices of early spinal processing and later spinal activity potentially shaped by supraspinal mechanisms. At the cortical level, early SEP components (N20, P25, N30, P45) predominantly reflect initial sensory encoding and sensorimotor integration, whereas later components (N60, P150) are more sensitive to attentional and cognitive influences [18,30,31]. Thus, SEPs and the NFR provide complementary measures of somatosensory and nociceptive processing during gentle touch.

The present work combined these measures to determine whether gentle touch with CT-optimal characteristics modulates sensory processing at the spinal level and/or at later supraspinal stages. It comprised an RIII experiment (Experiment 1) and two SEPs experiments (Experiments 2 and 3). Experiment 1 measured the NFR under four conditions: robot-delivered brushing, providing a standardized stimulus with controlled velocity and force; human-delivered stroking, adding the interpersonal context of real social touch; vibration, an active, predominantly Aβ-mediated control for generic tactile stimulation; and no touch, as baseline. We examined condition differences in RIII reflex amplitude and anticipated that gentle touch would attenuate it. Experiment 2 asked whether gentle touch modulates spinal and cortical SEPs relative to no touch, whether any such effect is specific to gentle touch or shared with other concurrent Aβ-mediated input, and whether it depends on the social nature of the touch. Median-nerve SEPs were recorded under the same four conditions as Experiment 1 to test whether gentle touch modulates spinal somatosensory processing, complementing the NFR findings. Because spinal responses are of low amplitude and difficult to resolve, Experiment 3 compared robot-delivered brushing with no touch using a high number of trials to increase sensitivity. We examined condition differences in early spinal processing (N13), late spinal processing potentially influenced by supraspinal mechanisms (P22), and cortical processing (N20, P25, N30, P45, N60, P150). We anticipated that gentle touch would attenuate N13 and increase P22, leave early cortical responses unaffected, and attenuate later cortical responses [32]. Through this approach, we aim to advance understanding of the neurophysiological mechanisms by which affective touch modulates somatosensation.

## 2. Methods

### 2.1. Participants

The same forty-four healthy volunteers (39 females, 5 males; 18–37 years; *M* = 21.25, *SD* = 3.53) took part in Experiments 1 and 2, on the same session (Session 1). All participants were students at the University of Minho and received academic credit for their participation. The sample size was established as a minimum of 35 participants, by balancing feasibility against prior electrospinography (ESG) studies and potential data quality issues. Regarding the Experiment 1, participants who did not end up with at least two trials of each condition were eliminated from the analysis, resulting in 35 participants with eligible NFR data (18–37 years; *M* = 21.3 *SD* = 3.83; 30 women, 5 men; 31 females, 4 males), similarly to other NFR studies [33,34]. Regarding the Experiment 2, as detailed in the Data processing section, 5 participants were subsequently excluded from the ESG SEPs analysis leaving a final sample of 39 participants. This final number is similar to the ESG study that informed our montage [17]. Experiments 1 and 2 were preregistered in the Open Science Framework (https://osf.io/s7rqd). Experiment 3 was conducted months after the other two, but before any data analyses. This last experiment was not preregistered, as the motivation behind this study is more exploratory and complementary to Experiment 2, and was motivated by a study published after the preregistration publication [16]. A separate sample of 17 participants (14 females, 3 males; 18 – 38 years; M=22.47, SD=4.74) took part in Experiment 3 in a different session. The sample size was informed by previous work using a comparable high-trial-count spinal SEP procedure [16].

Participants were required to have normal or corrected-to-normal vision and to be right-handed. To minimize sources of noise, they were advised to: arrive with loose, dry hair without any hair products; eat a light meal 1–3 hours before the session; abstain from alcohol for 24 hours; refrain from smoking and caffeine for at least 1 hour; and avoid wearing clothing covering the neck (e.g., turtlenecks). Exclusion criteria included chronic pain, acute pain on the day of testing, injuries to the arms, hands, or legs, psychiatric, neurological, or cardiovascular disorders, substance abuse, and diagnoses of major depressive disorder, anorexia nervosa, or autism spectrum disorder. All participants provided written informed consent before the experiment. The study protocol complied with the Declaration of Helsinki and was approved by the Ethics Committee for Social and Human Sciences of the University of Minho (ref. CEICSH 091/2023).

### 2.2. Self-report measures

Before the electrophysiological recordings of both sessions (Session 1: Experiment 1 and 2; Session 2: Experiment 3), participants completed a sociodemographic questionnaire. To further characterize the sample, after the recordings of Session 1, participants completed measures of touch experience, personality and interoception, which allowed us to examine whether any behavioral effects were associated to these individual differences, potentially influencing somatosensory processing. Touch-related experiences and attitudes were measured using the Portuguese version of the Touch Experiences and Attitudes Questionnaire [35,36], which comprises six subscales: Friends and family touch, Current intimate touch, Childhood touch, Attitude to self-care, Attitude to intimate touch, and Attitude to unfamiliar touch. Personality traits were assessed with the NEO Five-Factor Inventory [37,38], and interoceptive processes were assessed through the Body Perception Questionnaire (BPQ). The BPQ measures body awareness (sensitivity to internal bodily functions) and autonomic reactivity, which includes supradiaphragmatic and subdiaphragmatic symptom subscales.

### 2.3 Experimental Procedure

#### Tactile stimulation

CT-targeted brushing was applied to the left dorsal forearm by a robotic arm developed in-house. The dorsal forearm is supplied mainly by the C6–C8 dermatomes (C6 laterally, C7 centrally, C8 medially), whereas median-nerve afferents enter the cord through roots C6–T1, predominantly C6–C7, and generate N13 over the lower cervical cord [41,42]. The brushed skin and the stimulated nerve therefore overlap at C6, so CT input and the median-nerve volley can converge on the same segmental dorsal horn circuitry. The robotic device consisted of a cosmetic brush mounted on a robotic arm driven by a NEMA 17 stepper motor. Motor control was provided by a BIGTREETECH TMC2208 V3.0 stepper motor driver, enabling the delivery of smooth strokes at a CT-optimal velocity of approximately 3 cm/s while minimizing noise and vibration. The brush travelled along a 13-cm section of the forearm, with each stroke lasting approximately 3.5 s. To prevent visual feedback, the brush and motor were covered. For the human condition, the experimenter was trained to reproduce the same stimulation pattern by stroking the participant’s forearm (with the palm of the experimenter’s hand) at the same location and speed. The experiment included a sound cue delivered via headphones to the experimenter only. The trained experimenter used this cue to precisely time the start and end of each stroke during stimulation. Vibrotactile stimulation, targeting Aβ fibers conveyed primarily through the dorsal column-medial lemniscus pathway [43–45] was delivered using a vibrotactile device incorporating a linear resonant actuator (Minebea CLF-10WX01). The actuator was driven by a 100 Hz sinusoidal audio signal through a PAM8403 digital power amplifier board and housed within a 6 × 2.5 cm PLA plate, which was secured to the skin adjacent to the brushed area using adhesive tape.

Prior to Experiments 1 and 2, participants underwent a calibration phase to match the perceived intensity of vibration and brushing. Starting from zero intensity, participants gradually increased the vibration amplitude via a computer interface until it matched the subjective intensity of the brushing. This individualized vibration level was stored and used in the subsequent experiment.

In Experiments 1 and 2, after each condition block, participants rated the intensity of the sensation produced by the electrical stimuli according to a 0–100 numeric rating scale with verbal anchors that was placed above the stimulation screen (NRS:10 - “noticeable sensation”; 20 - “mildly painful sensation” (i.e., pain threshold); 30 - “very weak pain”; 40 - “weak pain”; 50 - “moderate pain”; 60 - “slightly strong pain”; 70 - “strong pain”; 80 - “very strong pain”; 90 - “nearly intolerable pain”; and 100 - “unbearable pain) and the pleasantness of the tactile stimulation (0 = very unpleasant; 100 = very pleasant).

#### Experiment 1 – NFR with four conditions

NFR were elicited from the biceps femoris of the left leg while electromyographic (EMG) activity was recorded. Participants were instructed to wear loose-fitting shorts to facilitate proper electrode placement. During the recordings, they sat in a reclining chair with the left leg supported at ∼60° hip and knee flexion. Electrical stimulation was delivered using the Digitimer DS7A stimulator. Electrodes were placed over the sural nerve at the retromalleolar pathway with conductive gel. Thresholding proceeded stepwise using a 0–100 NRS. Stimulation intensity was increased in 2 mA until participants reported moderate pain (∼50/100) on the NRS, after which intensity was decreased in 1 mA until participants rated less than 50, and then increased again until they rated 50. The experimental stimulation intensity was set at 1.5x this value. Reflex validity was confirmed online using ActiView, requiring a minimum EMG response of 20 µV (peak-to-peak) lasting ≥ 10 ms within 90–150 ms post-stimulus. If RIII wasn’t visible, the intensity would be increased. To ensure tolerability, the final stimulation level could not exceed 80 (“very strong pain”) on the NRS. Participants then completed four conditions: NFR with robot-delivered brushing, NFR with human stroking, NFR with vibrotactile stimulation, and NFR alone. Each condition comprised four trains of electrical pulses (16 trains total), with each train consisting of five rectangular pulses (0.5 ms duration, 100 Hz) with an inter-trial interval of 12 seconds. After each block, participants rated the intensity of the sensation produced by the electrical stimulation and the pleasantness of the tactile stimulation on a 0–100 (NRS) scale with verbal anchors (See Fig. 1A). Somatosensory stimulation was applied as described in the Tactile stimulation section.

**Fig 1.**
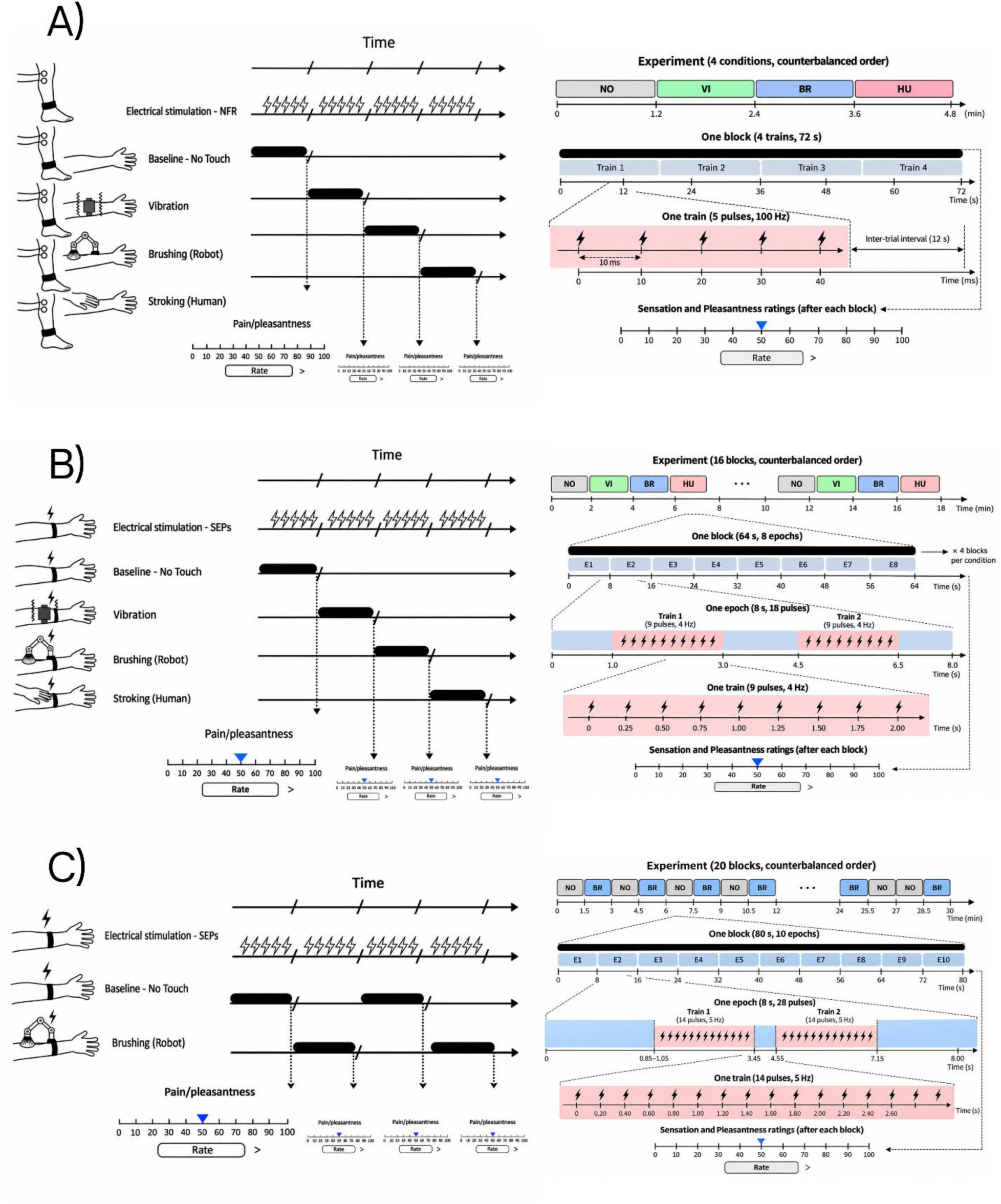
Visual depiction of the experimental procedures. A) Experiment 1: Electrical stimuli were applied to the sural nerve and NFR were recorded over bicep femoris during four experimental conditions: SEPs alone (no concomitant touch), SEPs + vibration (discriminative touch), SEPs + brushing (CT stimulation by a robot), and SEPs + Stroking (CT stimulation applied by a human). After each block, participants were asked to rate the sensation produced by the electrical stimulation and the pleasantness of the received sensory stimulation (vibration, brushing, or stroking) using a numeric rating scale (NRS). B) Experiment 2: Electrical stimuli were applied to the median nerve to record SEPs during the same four experimental conditions. Similarly, after each block, participants were asked to rate the sensation produced by the electrical stimulation and the pleasantness of the received sensory stimulation using an NRS. C) Experiment 3: Electrical stimuli were applied to the median nerve to record SEPs during two experimental conditions: SEPs alone (no concomitant touch) and SEPs + brushing (CT stimulation by a robot).

#### Electrical stimulation for SEPs

In Experiments 2 and 3, SEPs were elicited by stimulating the median nerve at the left wrist. Surface electrodes were positioned with the anode at the wrist crease and the cathode 2 cm proximal. Stimulation intensity was determined individually by gradually increasing the current in 2 mA steps until an involuntary twitch of ∼1 cm was observed in the median-innervated fingers. The experimenter then adjusted the intensity downward (1 mA) until the twitch disappeared and upward until it reappeared, repeating this procedure 2 times. Approximately 10 manual stimulations were delivered at the stimulus rate to confirm the final threshold (range 0.70-4.20 mA; *M* = 1.71 mA). Electrical stimulation was delivered as a rectangular constant-current pulse with a duration of 0.1 ms, with a Digitimer DS7A stimulator controlled by PsychoPy [46].

#### Experiment 2 - SEPs with four somatosensory conditions

Each participant received 2,304 electrical pulses across 16 counterbalanced blocks, with four blocks assigned to each condition and 576 pulses delivered per condition. Each block comprised eight consecutive 8 s stimulation epochs, resulting in 64 s of stimulation per block. Block order varied across participants and was fully counterbalanced across the sample. Within each epoch, 18 pulses were delivered in two trains of nine pulses, occurring from 1.0 to 3.0 s and from 4.5 to 6.5 s. Pulses within each train were presented at 4 Hz, corresponding to an interstimulus interval of 250 ms.

Four conditions were tested: electrical stimulation accompanied by robot-delivered brushing, human-delivered stroking, vibrotactile stimulation, or no concurrent touch. In the robot-brushing condition, the first electrical pulse was delivered approximately 1 s after initiation of the outward brush movement, whereas the return train began approximately 0.5 s after initiation of the return movement. The same parameters were used in the other two conditions, with electrical stimulation restricted to periods of brushing, stroking, or vibration (See Fig. 1B).

After each block, participants rated the intensity of the sensation produced by the electrical stimulation and the pleasantness of the tactile stimulation on a 0–100 (NRS) scale with verbal anchors. The experiment lasted approximately 18 minutes in total, excluding self-paced ratings and breaks.

#### Experiment 3 – SEPs with two somatosensory conditions

Each participant received 5,600 electrical pulses across 20 counterbalanced blocks, with 10 blocks assigned to each condition and 2,800 pulses delivered per condition. Each block comprised 10 consecutive 8 s stimulation epochs, resulting in approximately 80 s of stimulation per block. Within each epoch, 28 pulses were delivered in two trains of 14 pulses. Pulses within each train were presented at 5 Hz, corresponding to an interstimulus interval of 200 ms.

Two conditions were tested: electrical stimulation accompanied by robot-delivered brushing, and electrical stimulation without touch. Only two conditions were tested to allow a different analysis approach. Robot-delivered brushing was selected to control for socio-emotional salience of a human presence and no touch was selected as a baseline. In the tactile condition, each stimulation epoch encompassed the outward and return movements of the brush. The first electrical pulse was delivered approximately 0.85 to 1.05 s after the onset of the outward movement, and electrical stimulation was restricted to the outward and return brushing phases (See Fig. 1C). The experiment lasted approximately 30 minutes.

### 2.4 Electrophysiological Data Acquisition

Throughout Experiments 2 and 3, electrophysiological activity was continuously recorded using electroencephalography (EEG), electrospinography (ESG) and electrocardiography (ECG). The EEG was recorded with 47 active electrodes (Fp1, AF3, F1, F3, F5, FT7, FC5, FC3, FC1, C1, C3, C5, CP5, CP3, CP1, CPz, Pz, P1, P3, P5, POz, O1, Oz, Fpz, Fp2, AF4, AFz, Fz, F2, F4, F6, FT8, FC6, FC4, FC2, FCz, Cz, C2, C4, C6, CP6, CP4, CP2, P2, P4, P6, O2 – only cortical electrode names adhere to a standard system) placed over frontocentral and parietal scalp regions to capture cortical responses, including SEPs. The ESG was obtained using 16 electrodes surrounding one electrode positioned over the spinous process of the sixth cervical vertebra (C6) (17 electrodes total – S3, S4, S5, S6, S7, S8, S9, SC6, S11, S12, S13, S14, S15, S16, S17, S18, S19 – see Fig. 2C), and two additional reference electrodes – one under the seventh cervical vertebra (C7) and another on the glottis [17]. All physiological measures were recorded using a BioSemi ActiveTwo system sampled at 2048 Hz. The ECG was recorded using two electrodes attached bilaterally to the chest, approximately 2 cm below the clavicles. For the EMG used to record the NFR in Experiment 1, we used two Ag/AgCl electrodes positioned 5 to 10 cm above the popliteal fossa. All recordings took place in a sound-attenuated and light-controlled room.

**Fig 2.**
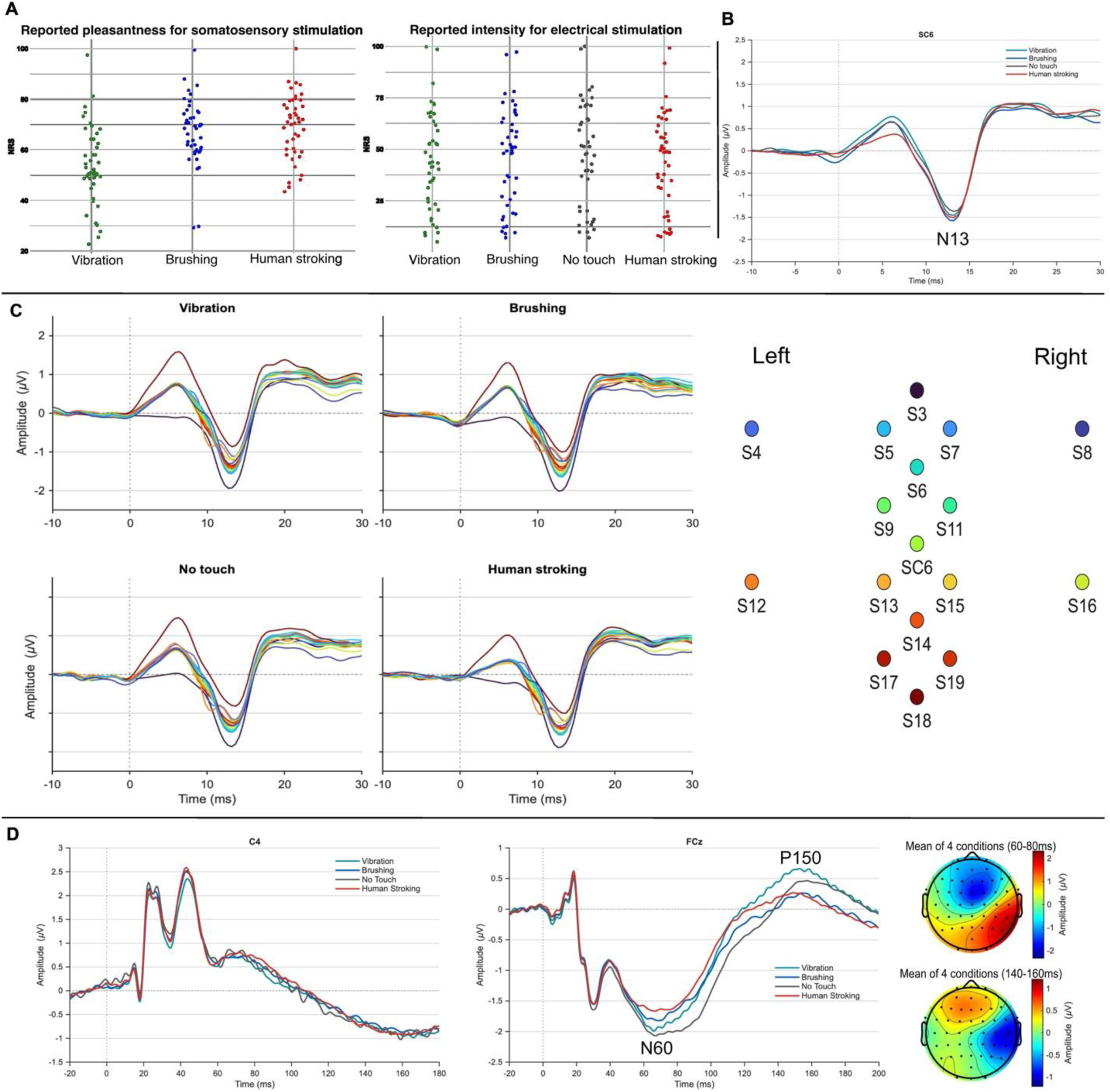
Results from Experiment 2. (A) Reported pleasantness for somatosensory stimulation (left) and reported intensity of electrical stimulation (right) for each condition on the Numeric Rating Scales (NRSs). (B) Spinal somatosensory evoked potentials for each condition recorded at electrode SC6; (C) Butterfly plots including all the spinal electrodes for each condition (Vibration, Brushing, No touch and Human stroking). On the right there is an illustration of the spatial layout of the electrodes and their associated color coding; (D) Cortical somatosensory evoked potentials recorded at electrode C4 (left) and FCz (right). The scalp topographies depict the mean activity across the four conditions in the 60-80 ms (top) and 140-160 ms (bottom) time windows.

**Fig 3.**
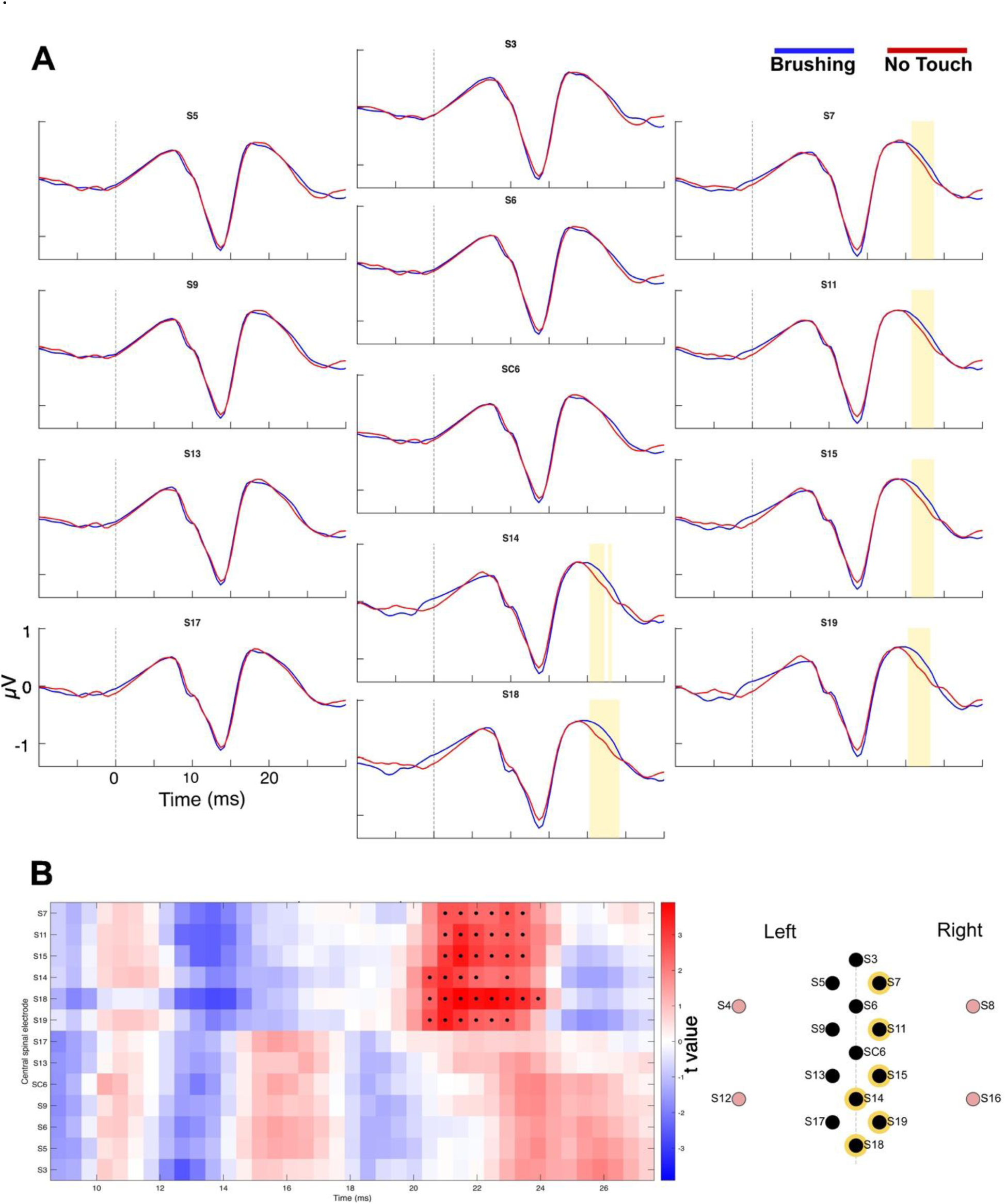
Spatiotemporal cluster-based permutation test in Experiment 3. (A) Grand-average spinal SEPs at the 13 analyzed electrodes, arranged to approximate their positions over the array. Blue: brushing; red: no touch. Yellow shading indicates the samples belonging to the significant cluster at around P22 latencies. (B) Left: dependent-samples t values (brushing minus no touch) across electrodes in the analysis window; warm colors indicate more positive amplitudes during brushing, and black dots indicate the samples forming the significant cluster. Right: electrode layout, with the four lateral electrodes excluded, shown in pink.

## 3. Data Analysis

### 3.1. Electrophysiological Processing

All EEG and ESG data were analyzed using the EEGLAB v2023.0 toolbox running under MATLAB R2023a. Raw data were preprocessed offline using custom scripts and EEGLAB functions. Recordings were imported using all EEG and ESG channels referenced to their average. Bipolar ECG, vertical EOG, and EMG signals were subsequently derived from their corresponding electrode pairs. To mitigate the large artifact generated by the electrical stimulator, samples falling within the 0 to 6 ms window after stimulus onset were replaced by a linear interpolation between the samples immediately preceding and following the artifact window. Artifact-contaminated segments—specifically those containing visible muscular activity or other non-neural transients— were removed following visual inspection.

For ESG data, a band-pass FIR filter (1–400 Hz) and the EEGLAB CleanLine procedure centered at 50 Hz were applied. Cardiac contamination of the ESG recordings was handled differently in the two experiments, following recent studies [16,17]. In Experiment 2, where fewer electrical stimuli were available for averaging, the spinal recordings were corrected using an R-peak–locked principal component analysis procedure, which has been shown to reduce the artefact level to less than 0.5% and to lead to a more than three-fold increase in SNR [47]. For this analysis, the 17 spinal channels, the additional neck channels positioned over the glottis and thoracic region, and the bipolar ECG channel were retained. R peaks were automatically detected from the ECG using the FMRIB QRS-detection algorithm. For each participant, the median R–R interval was calculated, and cardiac-artifact segments were extracted around each detected R peak using a window extending from 30% of the median R–R interval before the R peak to 50% after it. These R-locked segments were submitted to PCA separately for each spinal channel. A cardiac-artifact basis set was constructed from the mean R-locked waveform and the principal components that account for at least 90% of the variance, with a maximum of 5 PCA components. This basis set was fitted to each occurrence of the cardiac artifact by least-squares regression, and the fitted cardiac contribution was subtracted from the continuous ESG signal. After removing cardiac noise, spinal data were epoched from -40 ms before to 100 ms after electrical stimulus onset, with baseline correction applied using the –20 to –10 ms pre-stimulus interval. Epochs containing voltage exceeding ±150 µV across spinal channels were rejected.

In Experiment 3 (because it included a substantially larger number of electrical stimuli per condition), cardiac contamination was addressed using a more direct rejection-based approach. A different band-pass FIR filter (51–800 Hz) was applied and the baseline correction used the -80 to -5 ms interval. ECG R peaks were automatically detected, and SEP epochs containing an ECG R peak between −100 and +100 ms relative to the electrical stimulus were excluded from further analysis (total epochs: vibration: *M* = 524, range 404-575; robot brushing: *M* = 517.5, range 417-576; human stroking = 550.2, range 474-576; no touch: *M* = 520.5, range 428-574).

ESG waveforms for the N13 component were obtained by averaging all trials per condition. N13 was defined as the most negative peak between 10 and 16 ms post-stimulus. P22 was defined as the most positive peak between 18 and 25 ms. In Experiment 2, some electrodes were excluded from the analysis due to generally poor data quality in most participants and conditions, these electrodes were the ones placed closer to the borders of the neck patch: S3, S18, S4, S12, S8, S16, making them more prone to artifacts.

For EEG recordings, only data from Experiment 2 were included in the analyses, because the shorter inter-stimulus interval in Experiment 3 limited the reliable assessment of later cortical components. A band-pass FIR filter (1–200 Hz) and the EEGLAB CleanLine procedure centered at 50 Hz were applied. Cortical electrodes exhibiting excessive noise or poor contact were identified and reconstructed via spherical spline interpolation. Cortical data were epoched from – 100 ms before to 200 ms after electrical stimulus onset. For EEG data, ocular artifacts (e.g., blinks, lateral eye movements) were corrected using extended Infomax independent component analysis (ICA). The ICLabel plugin for EEGLAB [48] was used to classify each independent component (IC) into its seven categories. Components assigned a probability ≥ 0.80 of belonging to any of the artifactual classes (muscle, eye, heart, line noise, or channel noise) were automatically flagged and removed prior to further analysis.

EEG waveforms were obtained by averaging all trials per condition. Peak amplitude analysis targeted somatosensory evoked potential (SEP) components at two electrodes of interest. At electrode C4 (contralateral to the stimulated left hand), we measured N20 (most negative peak, 15–20 ms), P25 (most positive peak, 20–30 ms), N30 (most negative peak, 30–40 ms), and P45 (most positive peak, 35–50 ms). To complement these results, the significant differences over the same components in CP4 are also presented. At electrode FCz, we measured N60 (most negative peak, 40–100 ms) and P150 (most positive peak, 100–200 ms). The choice of the C4 electrode was due to its location over the contralateral S1 and because it is a broadly researched electrode in SEPs literature [18,30]. Regarding FCz, the choice was motivated by an established fronto-central maximum in SEPs at later stages of processing [18,30–32].

The EMG raw data were preprocessed offline using custom scripts and EEGLAB functions. A band-pass FIR filter (2–500 Hz) and a 50-Hz band-stop filter and its harmonics were applied. Artifact-contaminated segments were removed manually, i.e., sections that had either electric noise or contained a startle response in the first trial, producing substantially higher amplitudes that weren’t a result of the nociceptive stimulation. The RIII was quantified using z-score [22,49], determined by the difference between peak amplitude in the time window of 90 to150 ms poststimulation and mean baseline amplitude in the -60 to -10 ms before stimulus onset divided by the baseline standard deviation [33,50,51].

### 3.2. Statistical analyses

All analyses from Experiments 1 and 2 were conducted using Jamovi 2.6.26. One-way repeated measures ANOVAs were performed to analyse Numeric Rating Scale (NRS) scores, which assessed perceived intensity of the electrical stimulation and perceived pleasantness of the sensory stimuli, as well as electrophysiological data from Experiments 1 (EMG), and 2 (EEG and ESG) across four experimental conditions: SEPs without touch, SEPs concomitant with vibration, SEPs concomitant with brushing via robot and SEPs concomitant with human stroking.

For each ANOVA, we report partial eta squared (η²p) as the effect size. Where sphericity was violated, Greenhouse–Geisser corrected values are presented. Significant main effects were followed by post hoc pairwise comparisons using paired-samples *t* tests with Holm correction. To complement the frequentist analyses, Bayesian ANOVAs were conducted. We report Bayes factors for main effects and pairwise BF₁₀ for post-hoc comparisons, quantifying evidence in favor of the alternative hypothesis over the null, with BF₁₀ values between 1 and 3 interpreted as anecdotal evidence, between 3 and 10 as moderate evidence, and greater than 10 as strong evidence. In contrast to frequentist analyses, Bayesian statistics provides evidence for the null hypothesis, with values between 1/3 and 1 indicating anecdotal evidence, between 1/10 and 1/3 as moderate evidence, and less than 1/10 as strong evidence supporting the null hypothesis [52].

Associations between pain ratings and pleasantness ratings were examined using Kendall’s τ. As no significant correlations emerged, the results of these analyses are not reported in the results section. Correlations between NRS ratings and questionnaire measures were corrected for multiple comparisons using the false discovery rate (FDR), with adjusted pFDR values reported. As the questionnaire analyses were complementary to the study’s primary aims, their results are presented and discussed in the Supplementary Materials.

In Experiment 2, preregistered data eligibility included having 75% of the data which resulted in the exclusion of 5 participants, leading to a total of 39 participants with eligible spinal data. We retained 44 participants for the cortical data, as it is less sensitive to artifacts.

In Experiment 3 (epoch count: *M* = 3292; *SD* = 858.9), brushing and no-touch conditions were compared with a cluster-based permutation test in FieldTrip [53]. Dependent-samples t-statistics were calculated at each electrode and time point within a 20-millisecond window (from 8 to 28 ms post-stimulus). Samples exceeding the two-sided cluster-forming threshold (*p* < .05) were grouped into clusters according to temporal contiguity and spatial adjacency. Spatial neighbours were defined as electrodes separated by a Euclidean distance of 25 mm or less. Cluster-level significance was assessed using 5,000 within-subject label permutations and the maximum cluster-sum statistic. Family-wise error was controlled at α = .05 across both tails; consequently, clusters in each direction were evaluated against *p* < .025. The primary analysis included 13 central electrodes (S3, S5, S6, S9, S11, S13, S14, S15, S17, S18, S19, SC6, and S7), excluding the four lateral electrodes S4, S8, S12, and S16. To assess whether this exclusion influenced the findings, the analysis was repeated over the full 17-electrode array. Because the lateral contacts had no neighbours within 25 mm, a channel-specific neighbourhood radius was used for this sensitivity analysis: 40 mm for S4, S8, S12, and S16; 25 mm for all remaining electrodes. This connected the lateral contacts to their nearest electrodes and allowed them to contribute to clusters, although it resulted in a less homogeneous adjacency structure. A similar significant cluster was recovered using the 17-electrode array (S1 Fig).

## 4. Results

### 4.1. Experiment 1 – NFR with four conditions

#### 4.1.1. Behavioral data

##### Reported intensity for electrical stimulation

A one-way repeated measures ANOVA was conducted to examine the effect of condition in pain ratings in NFR, revealing a significant main effect (*F*(3, 102) = 9.58, *p* < .001, partial η² = 0.22, BF₁₀ =1520.78), and indicating strong evidence for the effect. Holm-corrected post-hoc comparisons indicated that, compared to the no-touch baseline, pain was significantly reduced during both human-delivered stroking (t(34) = 4.28, *p*< .001, BF₁₀ = 183.33), and robot-delivered brushing (t(34) = 3.20, *p* = .015, BF₁₀ = 12.24) with strong evidence for these differences. A significant difference was also observed between the human stroking and vibration condition (t(34) = 4.00, *p* = .002, BF₁₀ = 85.87) with strong evidence for this difference.

##### Reported pleasantness for sensory stimulation

There was also a significant main effect of stimulation type on pleasantness ratings (*F*(2, 68) = 14.4, *p* < .001, partial η² = 0.30, BF₁₀ = 5185.55) with strong evidence for this effect. Holm-corrected post-hoc comparisons indicated that human-delivered stroking was rated as significantly more pleasant than vibration (t(34) = 4.29, *p* < .001, BF₁₀ = 107.65). Similarly, robot-delivered brushing was also rated as significantly more pleasant than vibration (t(34) = 4.25, *p* < .001, BF₁₀ = 99.18). Both correlations show strong evidence for these differences. No significant difference was observed between human stroking and robot brushing.

To sum up, touch influenced pain ratings during NFR stimulation, although the effects differed across touch types. Both human-delivered stroking and robot-delivered brushing significantly reduced pain ratings relative to no touch, whereas vibration did not. Human stroking also produced lower pain ratings than vibration. Pleasantness followed a somewhat similar pattern: both stroking and brushing were rated as more pleasant than vibration, with no significant difference between them. Thus, although both forms of gentle touch were associated with greater pleasantness and reduced pain ratings, these effects were not entirely aligned, as human stroking showed a clearer advantage over vibration in pain reduction.

#### 4.1.2. Electrophysiological data

No significant effect of condition was found for RIII size (F(3, 102) = .79, *p* = .50, ηp² = .02, BF₁₀ = .10) with strong evidence for the null hypothesis (see Table 1).

**Table 1.** Descriptive Statistics RIII z-score.

|  | **Vibration** | **Brushing** | **No-Touch** | **Stroking** |
|---|---|---|---|---|
| **N** | 35 | 35 | 35 | 35 |
| **Mean** | 5.41 | 6.23 | 4.48 | 5.29 |
| **Median** | 2.04 | 1.22 | 1.58 | 1.16 |
| **SD** | 8.34 | 11.4 | 6.04 | 9.21 |
| **Minimum** | -0.158 | -0.207 | -0.224 | -0.158 |
| **Maximum** | 33.5 | 48.1 | 23.1 | 36.4 |

### 4.2. Experiment 2 – SEPs with four conditions

#### 4.2.1. Behavioral data

##### Reported intensity for electrical stimulation

Regarding the sensation/pain ratings, a one-way repeated-measures ANOVA revealed a significant main effect of stimulation type on sensation/pain ratings (*F*(2.48, 106.46) = 7.66, *p* < .001, partial η² = 0.15, BF₁₀ = 220), showing strong evidence for this effect (see Fig. 2A). Holm-corrected post hoc comparisons revealed that, relative to the no-touch baseline, sensation/pain ratings were significantly reduced during human-delivered stroking (t(43) = 4.24, *p* < .001, BF₁₀ = 206.57), providing strong evidence for a difference, and during vibration (t(43) = 3.33, *p* = .009, BF₁₀ = 17.48), providing moderate evidence for a difference. In contrast, robot-delivered brushing did not differ significantly from the no-touch baseline (t(43) = 2.32, *p* = .09, BF₁₀ = 1.79).

##### Reported pleasantness for sensory stimulation

A one-way repeated-measures ANOVA also revealed a significant main effect of stimulation type on pleasantness ratings (*F*(1.63, 69.89) = 24.5, *p* < .001, partial η² = 0.36, BF₁₀ = 4.65 × 10⁶), with strong evidence for this effect (see Fig. 2A). Holm-corrected post-hoc comparisons showed that both human-delivered stroking (t(43) = 5.77, *p* < .001, BF₁₀ = 20804.92) and robot-delivered brushing (t(43) = 5.02, *p* < .001, BF₁₀ = 2092) were rated as significantly more pleasant than vibration, both showing strong evidence for these differences.

To sum up, touch changed how the electrical stimulation was perceived, but not equally across touch types. Human-delivered stroking and vibration both reduced sensation/pain ratings relative to no touch, whereas robot-delivered brushing did not reliably do so, although the evidence for brushing was inconclusive. Pleasantness showed a different pattern: both stroking and brushing were rated as clearly more pleasant than vibration. Pleasantness and the reduction in sensation/pain ratings therefore did not go hand in hand. Vibration lowered ratings despite being the least pleasant, whereas brushing was pleasant without significantly lowering them. Only stroking showed both effects.

#### 4.2.2. Electrophysiological data

##### Spinal Activity

For all analyzed spinal electrodes, neither N13 nor P22 peak amplitudes revealed any significant differences between conditions. Repeated-measures ANOVAs (electrode: 4 levels) were conducted for the spinal recording sites (see Fig. 2B and C). No significant main effect of condition was found at any site (see Table 2). For N13, all p-values were above 0.23 and BF10 was between 0.07 and 0.17, providing moderate to strong evidence for an absence of condition differences. For P22, all p-values were above 0.09 and BF10 was between .15 and .45, indicating moderate to anecdotal evidence for an absence of condition differences.

**Table 2.** Results for spinal data in Experiment 2 – N13 and P22 – repeated-measures ANOVA.

|  | ***N13*** |  |  | ***P22*** |  |  |
|---|---|---|---|---|---|---|
| ***Electrode*** | ***F (3, 114 )*** | ***p-value*** | ***BF10*** | ***F (3, 114 )*** | ***p-value*** | ***BF10*** |
| **SC6** | 0.97 | 0.41 | .11 | 1.77 | 0.16 | .27 |
| **S19** | 0.96 | 0.42 | .10 | 1.43 | 0.24 | .18 |
| **S14** | 1.12 | 0.35 | .12 | 1.71 | 0.17 | .25 |
| **S17** | 1.26 | 0.29 | .15 | 1.26 | 0.29 | .15 |
| **S13** | 1.41 | 0.24 | .17 | 2.21 | 0.09 | .45 |
| **S15** | 0.69 | 0.56 | .07 | 1.79 | 0.15 | .27 |
| **S11** | 0.91 | 0.44 | .10 | 1.18 | 0.32 | .13 |
| **S9** | 0.95 | 0.42 | .10 | 1.74 | 0.16 | .25 |
| **S7** | 1.05 | 0.37 | .12 | 0.88 | 0.46 | .09 |
| **S5** | 0.77 | 0.51 | .08 | 1.27 | 0.29 | .15 |
| **S6** | 0.74 | 0.53 | .08 | 1.40 | 0.25 | .18 |

##### Cortical Activity

The earliest cortical responses were largely unaffected by the type of touch (see Table 3). Neither N20 (*F*(2.12, 91.26) = 2.64, *p* = .07, ηp² = .06, BF₁₀ = .67), N30 amplitude (*F*(1.90, 81.80) = 2.23, *p* = .12, ηp² = .05, BF₁₀ = .42) nor P45 (*F*(1.73, 74.43) = 1.74, *p* = .16, ηp² = .04, BF₁₀ = .23) differed across conditions, although the Bayes factors were only anecdotal and moderate, so these null results are inconclusive rather than evidence of absence. A clear effect appeared at P25 (*F*(3, 129) = 7.61, *p* < .001, ηp² = .15, BF₁₀ = 211.21), which was smaller during all three touch conditions than during no touch. This reduction was strong for vibration and modestly substantial for brushing and stroking. A similar effect on P25 was found at CP4 (F(3,129) = 3.67, *p* = .01, ηp² = .08, BF₁₀ = 2.24), with only anecdotal support.

**Table 3.**
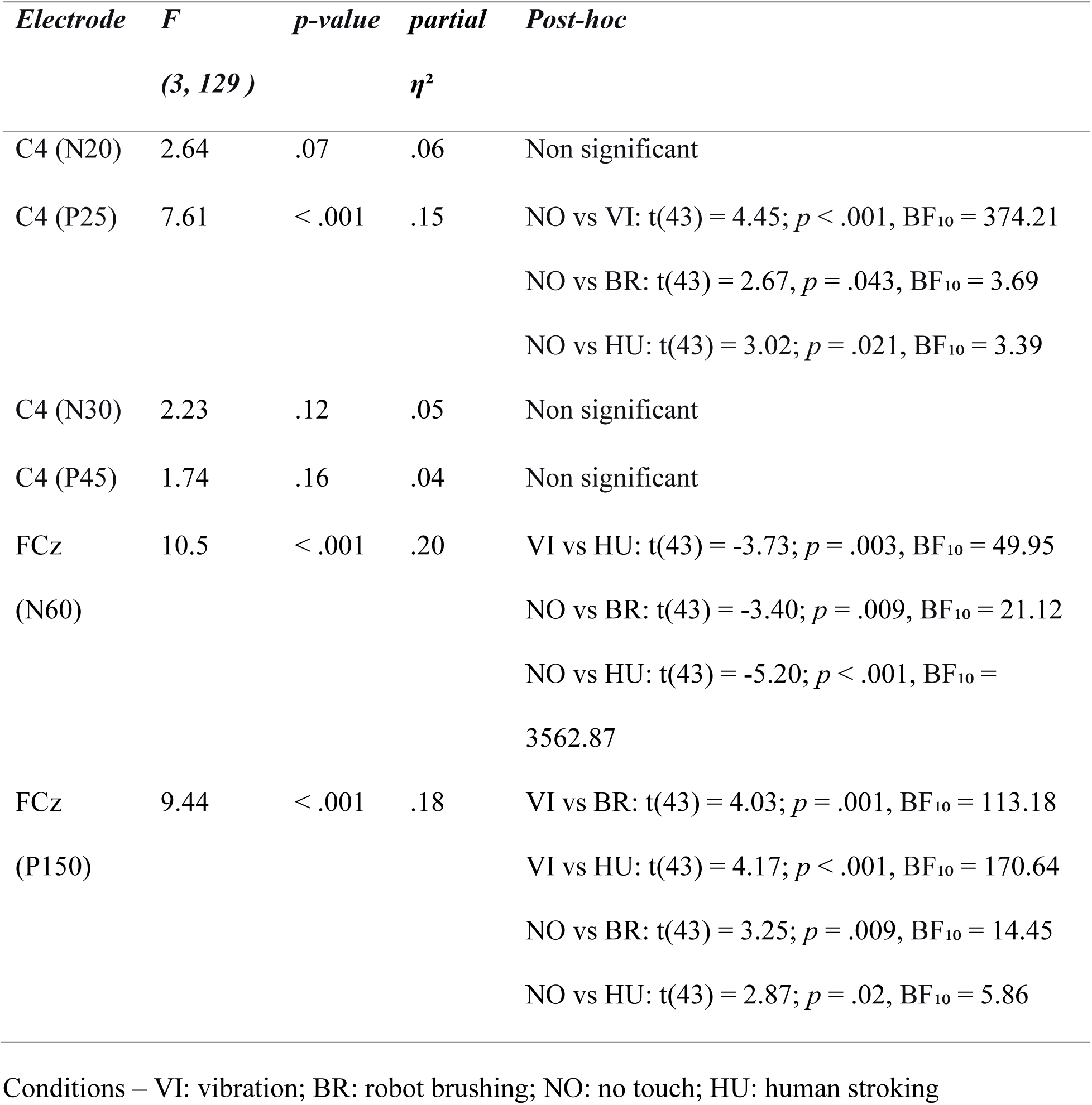
Results for cortical data in Experiment 2 - repeated-measures ANOVA.

| ***Electrode*** | ***F (3, 129 )*** | ***p-value*** | ***partial η²*** | ***Post-hoc*** |
|---|---|---|---|---|
| C4 (N20) | 2.64 | .07 | .06 | Non significant |
| C4 (P25) | 7.61 | < .001 | .15 | NO vs VI: t(43) = 4.45; *p* < .001, BF<sub>10</sub> = 374.21<br>NO vs BR: t(43) = 2.67, *p* = .043, BF<sub>10</sub> = 3.69<br>NO vs HU: t(43) = 3.02; *p* = .021, BF<sub>10</sub> = 3.39 |
| C4 (N30) | 2.23 | .12 | .05 | Non significant |
| C4 (P45) | 1.74 | .16 | .04 | Non significant |
| FCz (N60) | 10.5 | < .001 | .20 | VI vs HU: t(43) = -3.73; *p* = .003, BF<sub>10</sub> = 49.95<br>NO vs BR: t(43) = -3.40; *p* = .009, BF<sub>10</sub> = 21.12<br>NO vs HU: t(43) = -5.20; *p* < .001, BF<sub>10</sub> = 3562.87 |
| FCz (P150) | 9.44 | < .001 | .18 | VI vs BR: t(43) = 4.03; *p* = .001, BF<sub>10</sub> = 113.18<br>VI vs HU: t(43) = 4.17; *p* < .001, BF<sub>10</sub> = 170.64<br>NO vs BR: t(43) = 3.25; *p* = .009, BF<sub>10</sub> = 14.45<br>NO vs HU: t(43) = 2.87; *p* = .02, BF<sub>10</sub> = 5.86 |

Conditions – VI: vibration; BR: robot brushing; NO: no touch; HU: human stroking

The most robust condition effects emerged at frontocentral sites, at longer latencies (see Table 3 and Fig. 2E). N60 amplitude (*F*(3, 129) = 10.50, *p* < .001, ηp² = .20, BF₁₀ = 5071.45) was larger during no touch than during brushing and stroking, and larger during vibration than during stroking, with strong evidence for each comparison. P150 (F(3, 129) = 9.44, *p* < .001, ηp² = .18, BF₁₀ = 1657.82) showed the clearest pattern: strong evidence that it was smaller during both brushing and stroking when compared to vibration and strong to substancial evidence when compared to no touch.

Regarding our main hypothesis, gentle touch did not alter the earliest cortical components (N20, N30), but it modulated later frontocentral activity. The P25 reduction at C4 occurred for all touch conditions, including vibration, so it cannot be attributed specifically to gentle touch. In contrast, the FCz N60 and P150 were attenuated during brushing and stroking relative to no touch, and P150 was also attenuated relative to vibration in both gentle touch conditions. This pattern suggests that gentle touch may influence later stages of somatosensory processing.

### 4.2. Experiment 3 – SEPs with two conditions

#### Spinal Activity

A single significant spatiotemporal cluster was identified for the robot-delivered brushing > no touch contrast across the 13 central spinal electrodes and 39 time samples (two-sided FWER-corrected *p*=0.018). The cluster spanned 20.5 to 23.9 ms post-stimulus and included six electrodes: S19, S18, S14, S15, S11, and S7 (t-sum = 104.48). All remaining electrodes (n = 7) showed no significant clusters. Four lateral electrodes (S4, S8, S12, and S16) were excluded from this primary analysis because they had no neighbours within the 25 mm spatial-adjacency criterion. In the supplementary full-array analysis, including all 17 electrodes with an extended 40-mm neighbourhood radius for the four lateral contacts, a similarly significant cluster was recovered (See S1 Fig).

## 5. Discussion

The present study investigated whether gentle touch with CT-optimal characteristics modulates nociceptive and somatosensory processing already at the spinal level. We anticipated an attenuation in early spinal processing, an increase in later spinal processing and an attenuation in later cortical responses, using a threefold approach. Experiments 1 and 2 used a four-condition design (robot-delivered brushing, human-delivered stroking, vibrotactile stimulation, and a no-touch baseline) during both NFR and SEPs elicitation. Experiment 3 used a two-condition (robot-delivered brushing and no-touch) high-trial-count SEP design. Overall, the findings provided evidence that gentle touch did not modulate early spinal processing, as the NFR and N13 component did not differ across conditions, with Bayesian analyses even providing evidence for an absence of modulation. In contrast, gentle touch altered subjective ratings and cortical SEP components, particularly the frontocentral N60 and P150. In addition, Experiment 3 revealed an increase in a later spinal response around the latency of the P22 component, which is thought to reflect top-down influences on spinal processing. Together, these findings suggest that gentle touch modulates somatosensory processing beyond early segmental spinal activity, potentially through supraspinal mechanisms. Although both subcortical feedback loops and tonic corticospinal influences remain plausible, the temporal characteristics of the P22 response favor a subcortical contribution, potentially involving brainstem circuits, rather than modulation arising solely from local spinal processes.

The RIII reflex is primarily mediated by Aδ-fiber input and is widely used as an objective index of spinal nociceptive processing and central sensitization, with reduced thresholds commonly reported in chronic pain populations [50]. Although generated at the spinal level, its expression is also shaped by supraspinal influences, including threat and emotional context, demonstrating sensitivity to descending modulation [33,57]. The analysis of the NFR (RIII reflex z-scores) revealed no significant differences between brushing, stroking, vibration, and no-touch conditions, indicating no detectable changes in spinal nociceptive reflex excitability.

Behavioral findings contrasted with the absence of NFR modulation. Both human-delivered stroking and robot-delivered brushing significantly reduced pain ratings relative to no touch, despite producing no detectable changes in RIII amplitude. Both gentle touch conditions were also rated as more pleasant than vibration, with no difference between human and robotic touch. This dissociation between subjective pain and reflex responses suggests that gentle touch may influence pain through affective, attentional, or evaluative processes without measurable changes in spinal nociceptive reflex excitability. This interpretation is consistent with previous reports of reduced pain ratings without corresponding changes in RIII magnitude in healthy participants [21] and clinical populations [58].

In SEPs literature N13 component is considered a marker of early segmental somatosensory processing, reflecting postsynaptic activity within the deep dorsal horn, particularly laminae IV and V, following the arrival of large-diameter Aβ afferent input [24,25]. Its non-propagating spatial distribution and maximal amplitude over spinal segments corresponding to the stimulated nerve support its use as an index of segmental dorsal horn activity [17,26,27,54,55].

Across two experiments with complementary designs, we found no evidence that gentle touch modulated the N13. Previous studies showing N13 modulation have primarily used experimental nociceptive paradigms, reporting a decrease during heterotopic noxious conditioning stimulation induced by the cold pressor test [55], an increase following capsaicin-induced secondary hyperalgesia [26], and pain-model-dependent effects [56]. In contrast, our findings suggest that concurrent CT-optimal gentle touch does not measurably influence the dorsal horn populations contributing to the early Aβ-evoked N13 response to median nerve stimulation.

While the N13 is thought to reflect local segmental processing, the physiological origin of the later P22 component remains less certain. Previous work has proposed a contribution of afferent deporalization to this response [29], with recent evidence suggesting that its relatively late latency, broad rostrocaudal distribution, and sensitivity to attention are compatible with polysynaptic long-loop mechanisms involving supraspinal structures [16].

Here, although no effects emerged for N13, Experiment 3 revealed significantly more positive spinal activity during brushing between approximately 20 and 24 ms post-stimulus, corresponding to the expected P22 latency range. This effect was localized to the inferior right region, encompassing electrodes S14, S15, S18, and S19, that were contralateral to stimulation.

The timing of the P22 response provides important constraints on its potential supraspinal modulation. Because the P22 temporally overlaps with the cortical N20, any stimulus-evoked feedback contributing to this response would likely involve subcortical circuits, particularly at the brainstem level, rather than cortical processing. Indeed, the later cortical components modulated by gentle touch (N60 and P150) emerge too late to influence the P22 elicited by the same stimulus.

Alternatively, cortical influences may operate tonically, with ongoing touch establishing a modulatory state before each electrical pulse. Such modulation could involve attentional mechanisms, whereby directing attention toward touch and away from electrical stimulation enhances P22 amplitudes. Together with previous evidence of P22 sensitivity to attentional demands [16], these findings suggest that gentle touch may influence later spinal somatosensory processing through supraspinal mechanisms, while leaving the earlier N13 response unaffected. Notably, the P22 effect was not evident in Experiment 2. This discrepancy may reflect both the substantially lower number of trials per condition (576 vs. 2800), reducing signal-to-noise ratio, and differences in analytical approach. Detecting subtle, spatially distributed P22 effects may therefore require high trial numbers and sensitive methods such as cluster-based permutation testing.

At the cortical level, modulation was confined to later frontocentral responses, with no clear effects over somatosensory regions. Together with the absence of early spinal modulation, this pattern is consistent with a predominantly supraspinal contribution at higher-order stages of processing. In Experiment 2, both brushing and stroking attenuated frontocentral amplitudes relative to vibration and no-touch conditions. At N60, this attenuation was more consistent for human stroking (as there were no differences between vibration and brushing), whereas P150 was reduced during both gentle touch conditions compared to both vibration and no touch.

The N60, potentially generated within the supplementary motor area (SMA) and/or primary somatosensory cortex, has been linked to early sensory-attentional processing. Its attenuation may therefore reflect altered sensory-attentional processing during gentle touch. The P150 effect replicates our previous finding of reduced amplitudes during brushing relative to vibration and no touch [32]. Given the proposed involvement of this component in attentional allocation and later evaluative processing [59,60], its attenuation suggests that gentle touch preferentially influences later rather than initial cortical processing of somatosensory input.

Participants’ ratings showed a partly different pattern. Electrical stimulation was perceived as less intense during human stroking and vibration than during no touch, but not during robot-delivered brushing. Differences between human and robotic gentle touch have also been reported for pain modulation, and such effects appear to depend on the experimental paradigm [12,61].

Both human stroking and robot-delivered brushing were rated as significantly more pleasant than vibration consistent with the an established, but debatable [62], association between CT-targeted stimulation and pleasantness [6,63,64].

Importantly, whereas vibration also reduced perceived electrical stimulus intensity, only gentle touch attenuated the later frontocentral responses, particularly P150. This dissociation suggests that these cortical effects cannot be explained simply by reduced perceived stimulus intensity and may instead reflect the affective or evaluative characteristics of gentle touch [31].

## Limitations

Several methodological limitations should be considered. First, the sample was not sex-balanced. The observed P22 modulation may reflect general somatosensory activation rather than affective engagement, as there was no active tactile comparison. Regarding NFR, the lack of directly comparable studies raises uncertainty regarding the appropriateness of the stimulus count, warranting caution in interpretation. Moreover, because the NFR paradigm involved heterosegmental stimulation, its findings may not be directly comparable to those of the other two experiments.

The human-touch condition was also highly standardized, with the experimenter wearing headphones and following strict timing, which improved experimental control but may have reduced the naturalistic and socio-emotional qualities of interpersonal touch. Finally, although vibration was calibrated to match brushing intensity, human touch was not similarly matched, potentially contributing to differences in perceived stimulus intensity.

## Conclusion

The present findings indicate that gentle touch does not produce detectable modulation of early spinal somatosensory or nociceptive processing, as reflected by the N13 and NFR, while more consistent effects emerge at later stages. The P22 modulation observed in the high-trial-count experiment provides tentative evidence that gentle touch can influence later spinal processing, potentially through supraspinal mechanisms, although the physiological origin of this effect remains to be established.

At the cortical level, gentle touch attenuated the N60 and P150 SEP components relative to vibration and no touch, suggesting preferential modulation of later sensory-attentional and evaluative stages rather than initial cortical encoding. Importantly, these effects were observed for both robot- and human-delivered gentle touch, indicating that they cannot be attributed solely to the interpersonal nature of touch.

Overall, these findings argue against simple bottom-up inhibition within the spinal cord as the primary mechanism by which CT-targeted gentle touch shapes concurrent sensory and nociceptive processing. Instead, gentle touch appears to influence later stages of somatosensory processing at both spinal and cortical levels. Whether P22 modulation reflects a subcortical feedback loop or tonic corticospinal influences remains to be determined, although its temporal characteristics are consistent with a subcortical contribution.

By identifying the neural level at which CT-optimal gentle touch interacts with concurrent sensory input, these findings provide a mechanistic framework for understanding how socially and affectively meaningful touch shapes bodily experience without necessarily altering its initial somatosensory transmission.

## Data Availability Statement

Experiments 1 and 2 were preregistered on the Open Science Framework before the start of data acquisition and the preregistrations are openly available (https://osf.io/s7rqd). The data and code underlying the findings reported in this study, including electrophysiological recordings (EEG, ESG, and EMG) and associated behavioural data, are available through Zenodo. In line with the informed consent signed by participants, the data are divided into publicly available data and code (https://doi.org/10.5281/zenodo.24287670) and a restricted-access dataset (https://doi.org/10.5281/zenodo.24288118). Access to the restricted dataset can be requested through Zenodo and is granted to qualified researchers for scientific purposes, in line with the terms of the participants’ consent. Both datasets are organised according to the Brain Imaging Data Structure (BIDS) specification and include the relevant metadata and documentation.

## Supplementary Materials

### Psychological Measures Results

#### Reported intensity for electrical stimulation and questionnaires

No significant correlations were found between sensation/pain and pleasantness ratings, although some emerged in comparisons between pain and specific subscales of the TEAQ, BPQ, and NEO-FFI.

Regarding the comparison between sensation/pain and TEAQ, the only sub-scale with significant correlations was Attitude to self-care, that revealed a weak association with all sensation/pain ratings (stroking: τ = .25, *p* = .02, BF₁₀ = 3.45, *p*FDR = 0.18; brushing: τ = .26, *p* = .02, BF₁₀ = 3.52, *p*FDR = 0.18; vibration: τ = .26, *p* = .02, BF₁₀ = 3.80, *p*FDR = 0.18; no touch: τ = .24, *p* = .03, BF₁₀ = 2.70, *p*FDR = 0.18). However, none of the correlations survived FDR.

Similarly, the comparison between sensation/pain and NEO-FFI revealed significant differences between the Neuroticism dimension and all ratings (stroking: τ = .26, *p* = .02, BF₁₀ =3.83, *p*FDR = .08; brushing: τ = .26, *p* = .01, BF₁₀ = 4.11, *p*FDR = .08; vibration: τ = .27, *p* = .01, BF₁₀ = 4.93, pFDR = .08; no touch= τ = .27, *p* = .01, BF₁₀ =5.34, pFDR= .08) and Extraversion and both stroking and vibration (stroking: τ = .24, *p* = .03, BF₁₀ =2.55, *p*FDR = .08; brushing: τ = .20, *p* = .06, BF₁₀ = 1.20, *p*FDR = .15; vibration: τ = .25, *p* = .02, BF₁₀ = 2.93, *p*FDR = .08; no touch: τ = .21, *p* = .06, BF₁₀ = 1.28, *p*FDR = .15). All of these associations were weak and none survived FDR.

All sensation/pain ratings showed weak but statistically significant positive associations with BPQ-II scores (stroking: τ = .25, *p* = .02, BF₁₀ = 3.17, pFDR = .02; brushing: τ = .30, *p* = .004, BF₁₀ = 11.78, pFDR = .01; vibration: τ = .28, *p* = .008, BF₁₀ = 6.58, pFDR = .01; no-touch: τ = .28, *p* = .009, BF₁₀ = 5.92, pFDR = .01). The strongest evidence was observed for brushing, whereas the remaining conditions showed moderate evidence for an association. All associations survived FDR correction. When the BPQ-II was separated into supradiaphragmatic and subdiaphragmatic symptom subscales, sensation/pain ratings were associated only with supradiaphragmatic symptoms.

#### Reported pleasantness for sensory stimulation and questionnaires

Only a weak correlation emerged between pleasantness ratings during the vibration condition and the Friends and Family Touch subscale of the TEAQ (t = .26, *p* = .02, BF₁₀ = 3.70). This correlation did not survive FDR.

## Discussion

Regarding associations between NRS ratings and questionnaire measures, the most consistent findings emerged for the BPQ-II supradiaphragmatic subscale. Participants reporting more supradiaphragmatic autonomic symptoms also rated the electrical stimulation as more intense across all experimental conditions. BPQ-II reflects heightened awareness and sensitivity to bodily sensations, the supradiaphragmatic subscale focuses on autonomically innervated organs above the diaphragm, being associated with social engagement and resting state vs “fight or flight” responses. Our results suggest that individual differences in the reporting or perception of bodily symptoms are associated with the perceived intensity of somatosensory stimulation, independent of the concurrent tactile condition. This finding is consistent with previous work linking enhanced autonomic sensitivity and pain sensitivity [65–67].

**S1 Fig.**
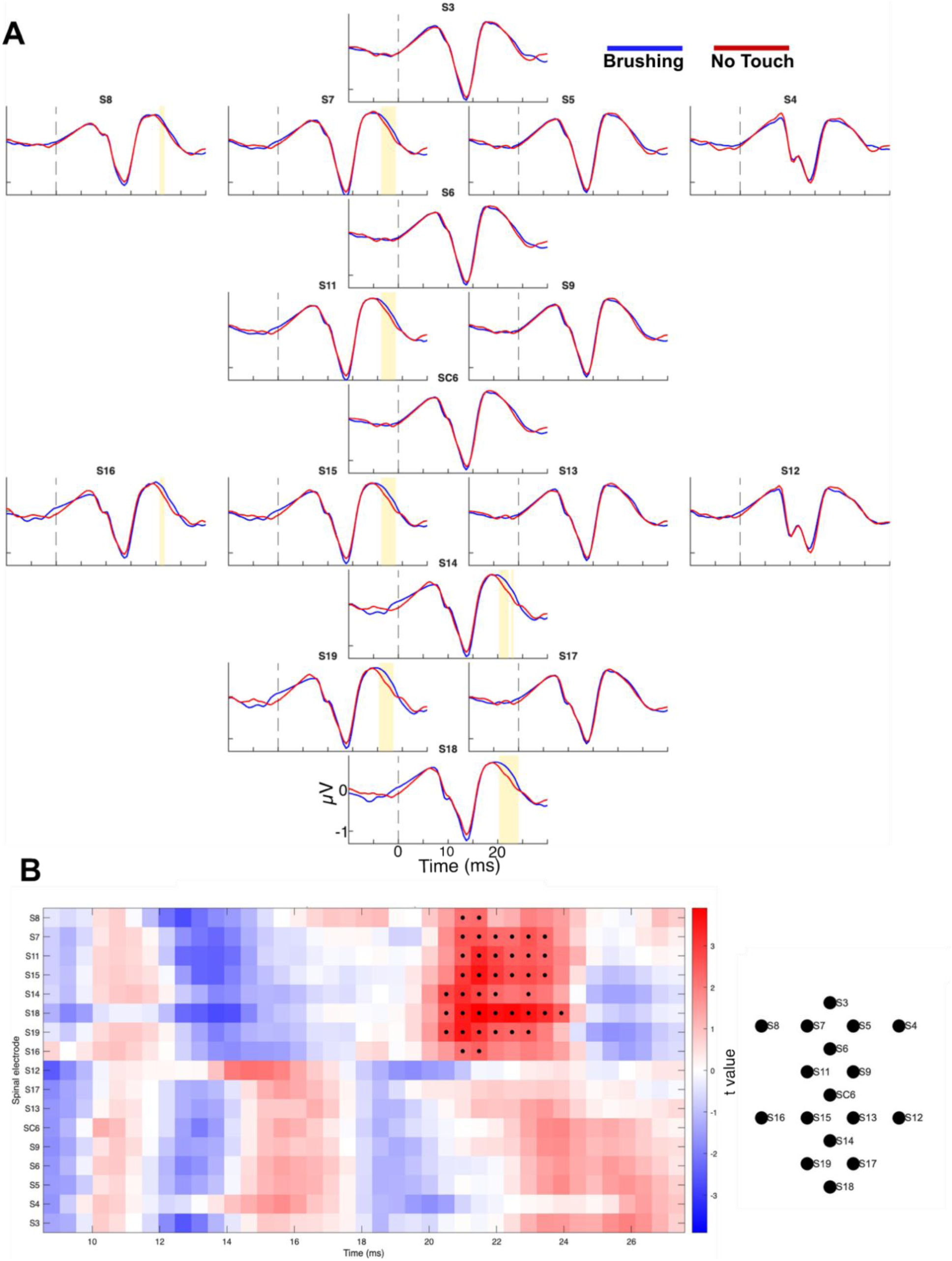

### Spatiotemporal cluster-based permutation test in Experiment 2, including the 4 lateral electrodes

(A) Grand-average spinal SEPs at the 17 analyzed electrodes, arranged to approximate their positions over the array. Blue: brushing; red: no touch. Yellow shading indicates the samples belonging to the significant cluster at around P22 latencies. (B) Left: dependent-samples t values (brushing minus no touch) across electrodes in the analysis window; warm colors indicate more positive amplitudes during brushing, and black dots indicate the samples forming the significant cluster. Right: electrode layout.

